# Sotagliflozin, a dual SGLT1/2 inhibitor, improves cardiac outcomes in a mouse model of early heart failure without diabetes

**DOI:** 10.1101/2020.08.15.252593

**Authors:** Sophia L Young, Lydia Ryan, Thomas P Mullins, Melanie Flint, Sarah E Steane, Sarah L Walton, Helle Bielefeldt-Ohmann, David A Carter, Melissa E Reichelt, Linda A Gallo

## Abstract

**Aims:** Selective SGLT2 inhibition reduces the risk of worsening heart failure and cardiovascular death in patients with existing heart failure, irrespective of diabetic status. We aimed to investigate the effects of dual SGLT1/2 inhibition, using sotagliflozin, on cardiac outcomes in non-diabetic and diabetic mice with cardiac pressure overload.

**Methods and Results:** Five-week old male C57BL/6J mice were randomized to receive a high fat diet (HFD; 60% of calories from fat) to induce diabetes or remain on normal diet (ND) for 12 weeks. Transverse aortic constriction (TAC) was then employed to induce cardiac pressure-overload (50% increase in right:left carotid pressure versus sham surgery), resulting in features representative of heart failure with preserved ejection fraction. At five weeks into the dietary protocol, mice were treated for seven weeks by oral gavage once daily with sotagliflozin (10mg/kg body weight) or vehicle (0.1% tween 80). In ND non-diabetic mice, treatment with sotagliflozin attenuated cardiac hypertrophy and histological markers of cardiac fibrosis induced by TAC. These benefits were associated with profound diuresis and glucosuria, without shifts toward whole-body fatty acid utilisation nor increased cardiac ketolysis. In HFD diabetic mice, sotagliflozin did not attenuate cardiac injury induced by TAC. HFD mice had vacuolation of proximal tubular cells, associated with less profound diuresis and glucosuria, which may have compromised drug action and subsequent cardio-protection.

**Conclusion:** We demonstrate the utility of dual SGLT1/2 inhibition in treating heart failure risk factors in the non-diabetic state. Its efficacy in high fat-induced diabetes with proximal tubular damage requires further study.

## Introduction

Sodium-dependent glucose transporter (SGLT)-2 inhibitors have emerged as promising anti-diabetic agents, exerting rapid cardiovascular benefits beyond what would be expected from adequate glycaemic control (1-3). Irrespective of diabetic status, baseline HbA_1c_, or degree of glucose lowering, the SGLT2 inhibitor, dapagliflozin, reduced the risk of worsening heart failure or cardiovascular death in patients with existing heart failure (4, 5). This led to the recent FDA approval for dapagliflozin in the treatment of heart failure in adults with and without type 2 diabetes mellitus (DM).

SGLT2, located in the early kidney proximal tubule, is responsible for up to 97% of renal glucose reabsorption, while SGLT1, in the late proximal tubule, removes any remaining luminal glucose (6). In patients with impaired renal function, selective SGLT2 inhibitors have reduced efficacy and, in all type 2 DM patients, modest effects on blood glucose lowering are observed: complete SGLT2 inhibition induces maximal glucosuria of only 40-60% due to compensatory upregulation of SGLT1-mediated kidney glucose reabsorption (6, 7). SGLT1 is also the main transporter for intestinal glucose absorption (6). Consequently, co-inhibition of SGLT1 and SGLT2 is being explored to provide greater blood glucose lowering effects and, potentially, cardiovascular benefits.

Sotagliflozin is a dual SGLT1/2 inhibitor, with approximately 20-fold selectivity for SGLT2 over SGLT1 (6). Clinical trials in type 2 DM patients have demonstrated a favourable safety profile with sotagliflozin, as well as reductions in HbA_1c_, body weight, and blood pressure (8, 9), and an ongoing phase 3 trial is examining the effects of sotagliflozin on cardio-renal outcomes (NCT03315143). It is likely that dual SGLT1/2 inhibitors will offer vascular benefits analogous, or perhaps even superior, to selective SGLT2 inhibitors. However, unlike SGLT2, SGLT1 is also expressed in the heart; most likely localised to heart capillaries (10-12) and hence may also have a direct cardiac effect.

Increased myocardial SGLT1 expression has been reported in various disease states in humans (end-stage cardiomyopathy and type 2 DM) and mice (myocardial ischaemia, type 2 DM, and glycogen storage cardiomyopathy) (11, 13). Furthermore, transgenic cardiomyocyte-specific SGLT1 knockdown attenuated disease features in mice with glycogen storage cardiomyopathy (14). In contrast, in non-diabetic isolated murine hearts subjected to ischemia reperfusion injury, non-selective SGLT inhibition with phlorizin impaired functional recovery, associated with decreased glucose uptake and ATP content (10), suggesting a putative role in heart function. The functional role of myocardial SGLT1 remains largely unclear. Accordingly, we aimed to investigate the effects of dual SGLT1/2 inhibition, using sotagliflozin, on cardiac outcomes in a mouse model of cardiac pressure overload in the presence and absence of type 2 DM (15).

## Methods

### Experimental protocol

All procedures were performed in accordance with guidelines from The University of Queensland Animal Ethics Committee and the National Health and Medical Research Council of Australia (#334/16). Four-week old male C57BL/6J mice were purchased (Australian Resource Centre, Murdoch, Australia) and housed in an environmentally controlled room (constant temperature 22°C), with a 12 h light-dark cycle (lights on at 6:00 h and off at 18:00 h) and access to standard chow and RO water *ad libitum*. At five weeks of age, mice were randomized to receive a high fat diet (HFD) or remain on normal diet (ND) for 12 weeks. The HFD was 21.7 MJ/kg; 60% of calories from fat, 20% from protein, and 20% from carbohydrate (13-092, Specialty Feeds, Glen Forrest, Australia) and the ND was 14 MJ/kg; 12% of calories from fat, 23% from protein, and 65% from carbohydrate. At one week into the dietary protocol, transverse aortic constriction (TAC) was employed to induce cardiac pressure-overload (15). Here, mice were anesthetized with a mixture of ketamine and xylazine (80/8 mg/kg) and a blunt 26-gauge needle was positioned over the transverse aorta. The suture was tightened on the shaft of the needle to standardise the degree of aortic restriction and the needle immediately removed. Sham-operated mice were also included whereby the same procedures were carried out except for aortic constriction. At five weeks into the dietary protocol, mice were treated for seven weeks by oral gavage once daily with either sotagliflozin (SOTA, 10mg/kg body weight) or vehicle (0.1% tween 80), resulting in eight groups (n=6-7/group). Body weight and fasting blood glucose levels were monitored throughout the study. At study completion (12 weeks on high fat diet/11 weeks post-TAC), mice were anesthetized with a mixture of ketamine and xylazine (80/8 mg/kg). The carotid pressure differential was assessed using pressure catheters inserted into left and right carotid arteries and simultaneously recorded for five min (SPR1000; Millar Inc., Houston, USA).

Hearts and kidneys were excised and snap-frozen in liquid nitrogen or fixed in 10% neutral-buffered formalin for molecular and histological analyses, respectively. All physiological measurements described below were carried out in the last two weeks of study.

### Glucose tolerance and body composition

An oral glucose tolerance test (OGTT) was performed following a 5-6 h fast between 08:00-14:00 h using 50% w/v D-glucose solution (2 g/kg body weight) and a SensoCard glucometer (15). Body fat content was measured in conscious mice using nuclear magnetic resonance (Bruker’s Minispec MQ10, Houston, USA).

### Echocardiography

Transthoracic echocardiography (M-mode) was performed in a blinded fashion by an experienced veterinarian (M.F.) using a Philips iE33 ultrasound machine with a 15 MHz linear transducer (Philips, Amsterdam, Netherlands) under 1.8% inhaled isoflurane anesthesia, as previously described (15). All data are the average of at least three consecutive beats.

### Kidney function

Mice were weighed and placed individually into metabolic cages for 24 h urine collection (15). Mice were acclimatized to the metabolic cages by housing for a short period of daylight prior to the 24 h collection. An ELISA kit was used for the measurement of urinary albumin (Bethyl Laboratories, Montgomery, USA).

### Indirect calorimetry

An automated open-circuit indirect calorimetry system (TSE Systems, Bad Homburg, Germany) was used for the measurement of food and drink patterns, and energy balance, as previously described (15). This system comprised one environmental chamber, which allowed for individual monitoring of 12 mice in parallel with high resolution and precision. Mice were acclimatized to the new housing conditions for seven days, followed by continuous recording for three consecutive days. Data were recorded for each animal every hour. To calculate day- and night-time averages, a single value per animal was used (the average of three 12-hr cycles).

### Heart and kidney histology

Paraffin sections (4 µm) were used for the blinded assessment of heart (cross-sectional plane) and kidney injury. All sections were visualized using an Aperio Slide Scanner. Myocyte size (average of 50 cross-sectional myocytes per animal) was determined in the short axis plane in the left ventricle following Picrosirius red staining. A descriptive histopathology assessment was performed by an expert pathologist (H.B.O.) using Masson’s Trichrome and Picrosirius red staining for hearts and Masson’s Trichrome, PAS, and H&E staining for kidneys.

### Heart and kidney real-time qPCR

Total RNA was extracted from heart ventricles and kidney cortex using Trizol. cDNA was synthesized using iScript cDNA synthesis and real-time qPCR performed using pre-designed Taqman Gene Expression Assays for *Myh7* (Mm00600555_m1), *Nppa* (Mm01255747_g1), *Fibronectin* (Mm01256744_m1), *ColIVα1* (Mm01210125_m1), *Slc5a1* (Mm00451203_m1), *Slc5a2* (Mm00453831_m1), *Mct2* (Mm00441442_m1), and *Bdh1* (Mm00558330_m1) (Life Technologies, Mulgrave, VIC, Australia). Relative gene expression was quantified using the comparative threshold cycle (ΔΔCt) with 18S rRNA (Life Technologies) as the endogenous multiplexed control.

### Statistical analyses

Data were analyzed by three-way ANOVA (SPSS, HFD x TAC x SOTA) or two-way ANOVA (GraphPad Prism 7, TAC x SOTA within each diet group). Upon significant interaction, data were split where appropriate and analyzed by two-way ANOVA or Tukey’s multiple comparisons test. All data were tested for normality using the Shapiro-Wilk test and log transformed for statistical analyses, where required. Data are expressed as means ± SD for normally distributed data or median ± IQR for not normally distributed data. *P* < 0.05 was considered significantly different.

## Results

HFD induced a diabetic state, reflected by elevated fasting blood glucose levels, impaired glucose tolerance (*Figure 1A-D*), and fasting and glucose-stimulated hyperinsulinaemia (*Figure 1E-F*). There was no effect of HFD on total body mass, but fat mass was significantly increased compared with mice on ND (*Figure 1G-H*). TAC surgery successfully induced cardiac pressure overload in both ND and HFD mice (50% increase in right-to-left carotid artery pressure, *P*<0.05, data not shown), associated with left ventricular hypertrophic remodeling (*Figure 2A-D* and *Table 1-2*). Histological assessment of Trichrome-stained slides revealed a greater proportion of TAC mice with a degree of pathology including fibrosis, myocyte hypertrophy, and/or scattered leukocyte infiltration compared with SHAM (*Figure 2C*). Fractional shortening and ejection fraction were preserved in this model of cardiac pressure overload (*Table 1-2*). TAC surgery in HFD, but not ND, mice altered cardiac expression levels of genes involved in ketolysis, with decreased ketone body transporter, *Mct2*, and increased ketolytic enzyme, *Bdh1* (*Figure 2D*). In ND mice, TAC surgery reduced total energy and water intake, associated with reduced energy expenditure (*Figure 4A-B, Figure 5A*). This effect was not seen in HFD mice.

**Table 1.** Echocardiography parameters in ND mice.

|  | SHAM |  | TAC |  | *P value* |  |
| --- | --- | --- | --- | --- | --- | --- |
|  | VEH | SOTA | VEH | SOTA | *TAC* | *SOTA* |
| LVIDd (mm) | 3.626 ± 0.280 | 3.552 ± 0.392 | 3.351 ± 0.500 | 3.781 ± 0.328 | NS | NS |
| LVAWd (mm) | 0.844 ± 0.089 | 0.883 ± 0.166 | 1.102 ± 0.200 | 0.899 ± 0.146 | <0.05 | NS |
| LVPWd (mm) | 0.987 ± 0.230 | 0.899 ± 0.105 | 1.546 ± 0.528 | 1.059 ± 0.154 | <0.01 | <0.05 |
| LVIDs (mm) | 2.574 ± 0.463 | 2.229 ± 0.467 | 2.195 ± 0.435 | 2.715 ± 0.454 | NS | NS |
| LVAWs (mm) | 1.013 ± 0.139 | 1.199 ± 0.145 | 1.365 ± 0.243* | 1.172 ± 0.088 | <0.05 | NS |
| LVPWs (mm) | 1.284 ± 0.319 | 1.381 ± 0.233 | 1.960 ± 0.551* | 1.373 ± 0.277# | <0.05 | NS |
| FS (%) | 29.41 ± 9.18 | 37.79 ± 7.94 | 34.83 ± 6.71 | 28.54 ± 7.73 | NS | NS |
| EF (%) | 63.26 ± 12.07 | 74.88 ± 8.58 | 71.43 ± 8.04 | 62.38 ± 11.03 | NS | NS |

Data are means ± SD (*n*=6-7/group). \* *P* < 0.05 vs SHAM counterpart, # *P* < 0.05 vs VEH counterpart by Tukey's *post hoc* following significant interaction by two-way ANOVA. NS=not significant; LVIDd/s=left ventricular internal diameter in diastole/systole; LVAWd/s=left ventricular anterior wall thickness in diastole/systole; LVPWd/s=left ventricular posterior wall thickness in diastole/systole; FS=fractional shortening; EF=ejection fraction.

**Table 2.** Echocardiography parameters in HFD mice.

|  | SHAM |  | TAC |  | *P value* |  |
| --- | --- | --- | --- | --- | --- | --- |
|  | VEH | SOTA | VEH | SOTA | *TAC* | *SOTA* |
| LVIDd (mm) | 3.574 ± 0.386 | 3.498 ± 0.334 | 3.380 ± 0.156 | 3.653 ± 0.203 | NS | NS |
| LVAWd (mm) | 0.926 ± 0.104 | 0.868 ± 0.091 | 0.949 ± 0.079 | 1.061 ± 0.128 | <0.05 | NS |
| LVPWd (mm) | 1.043 ± 0.230 | 1.097 ± 0.251 | 1.150 ± 0.205 | 1.201 ± 0.215 | NS | NS |
| LVIDs (mm) | 2.410 ± 0.542 | 2.537 ± 0.337 | 2.311 ± 0.202 | 2.656 ± 0.238 | NS | NS |
| LVAWs (mm) | 1.136 ± 0.158 | 1.065 ± 0.074 | 1.188 ± 0.114 | 1.235 ± 0.131 | =0.054 | NS |
| LVPWs (mm) | 1.386 ± 0.339 | 1.413 ± 0.200 | 1.431 ± 0.236 | 1.535 ± 0.202 | NS | NS |
| FS (%) | 32.83 ± 11.17 | 27.41 ± 6.96 | 31.69 ± 3.02 | 27.27 ± 5.54 | NS | NS |
| EF (%) | 67.50 ± 15.07 | 60.71 ± 11.21 | 67.91 ± 4.35 | 60.8 ± 8.51 | NS | NS |

Data are means ± SD (*n*=5-6/group). No significant interactions observed by two-way ANOVA. NS=not significant; LVIDd/s=left ventricular internal diameter in diastole/systole; LVAWd/s=left ventricular anterior wall thickness in diastole/systole; LVPWd/s=left ventricular posterior wall thickness in diastole/systole; FS=fractional shortening; EF=ejection fraction.

**Figure 1.**
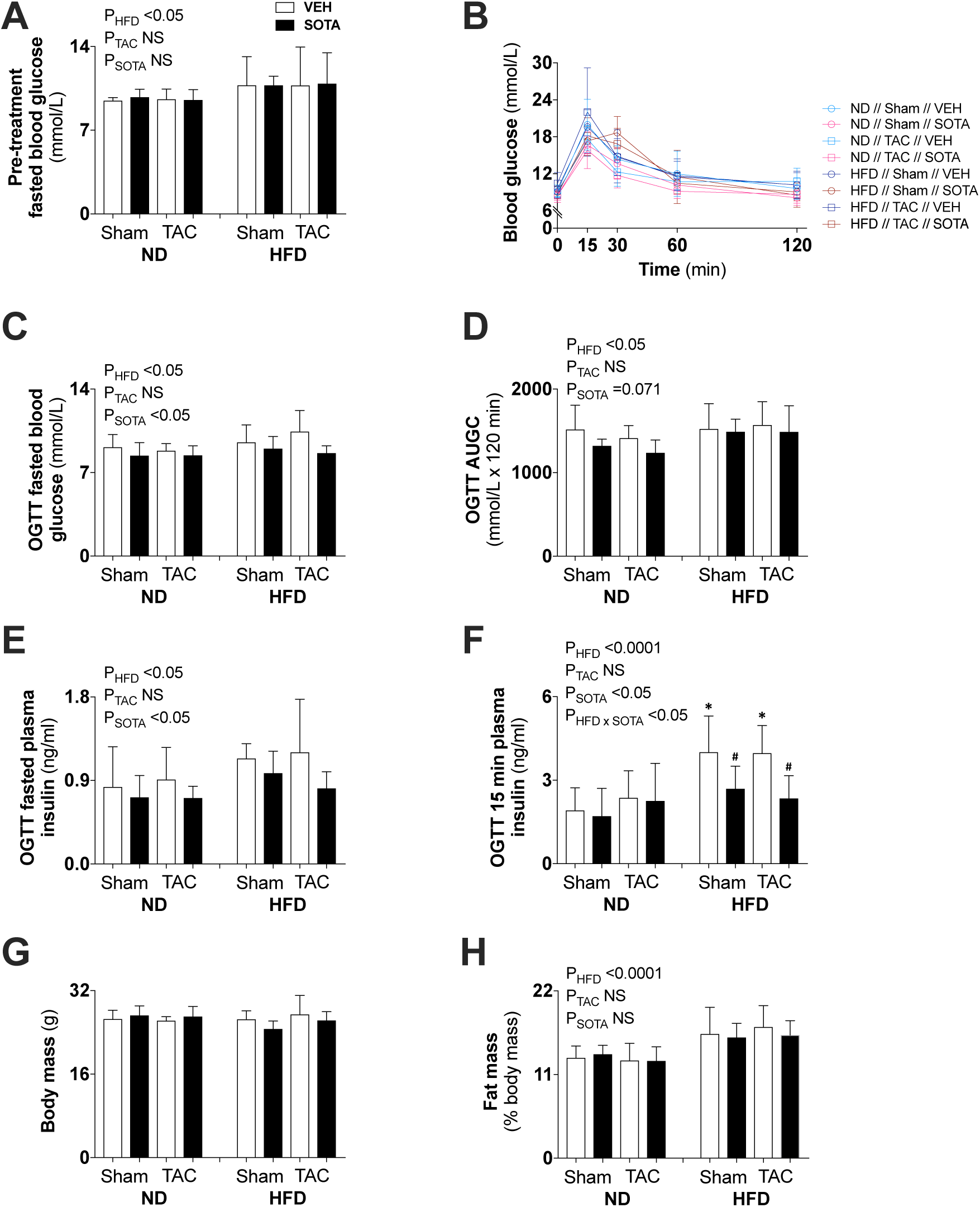
Glucose and insulin levels, and body and fat mass in non-diabetic and diabetic mice with cardiac pressure overload treated with sotagliflozin. *(A)* baseline (pre-treatment) fasted blood glucose levels, (*B-F*) oral glucose tolerance test (OGTT) blood glucose profile, fasted blood glucose levels, area under the glucose curve (AUGC), fasted plasma insulin levels, and 15-minute plasma insulin levels, (*G*) body mass, and (*H*) fat mass relative to body mass. Data are means ± SD (*n*=5-7/group). * *P* < 0.01 *vs* ND counterpart, # *P* < 0.01 *vs* VEH counterpart following significant interaction by three-way ANOVA. Interaction p-values indicated only when significant (or trend). NS=not significant.

**Figure 2.**
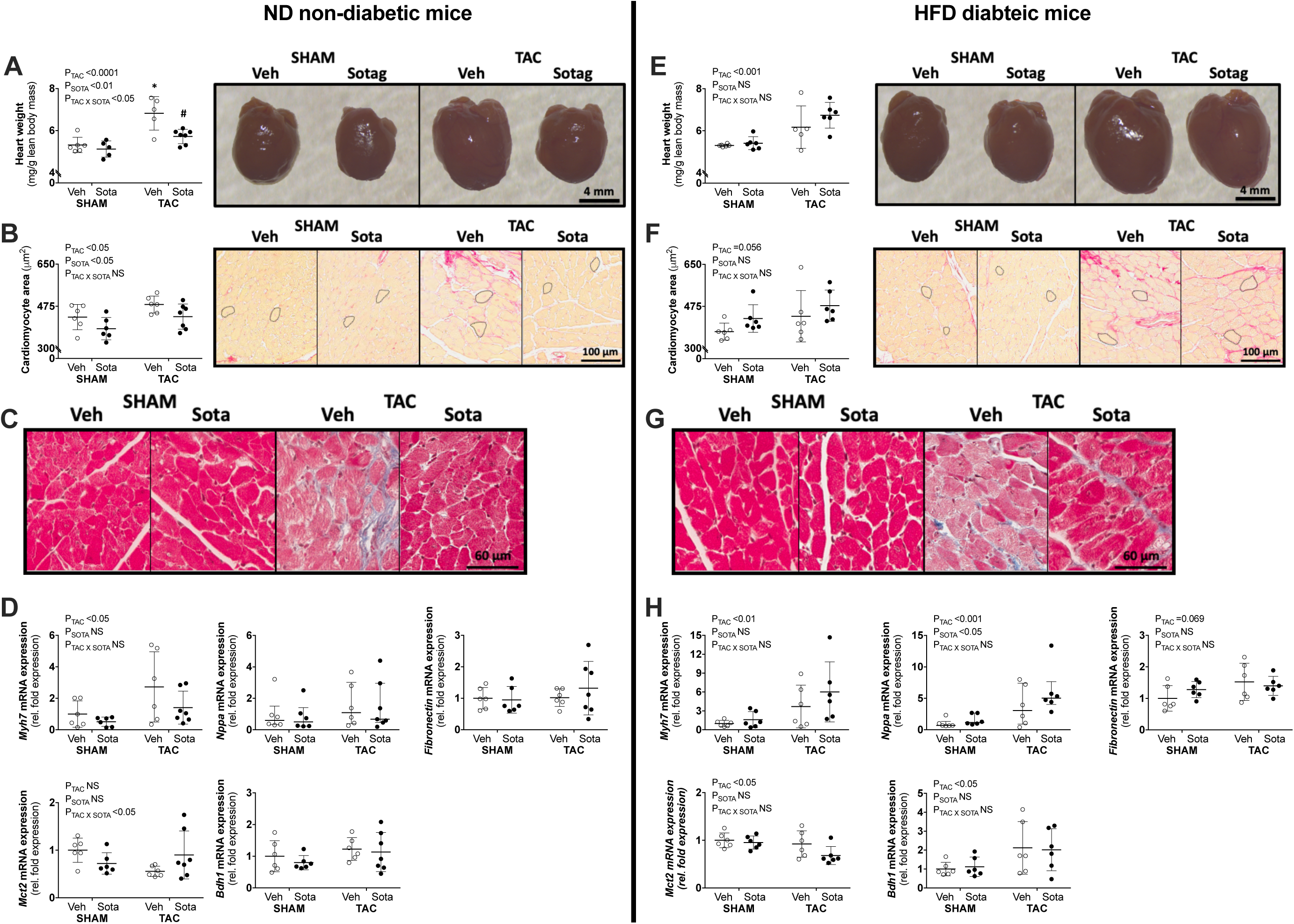
Effect of sotagliflozin on heart size, cardiomyocyte area, cardiac fibrosis, and cardiac gene expression in non-diabetic and diabetic mice with cardiac pressure overload. *(A, E)* heart weight relative to lean body mass, (*B, F*) cardiomyocyte area, (*C, G*) Trichrome-stained representatives used in descriptive histological assessments by expert pathologist (H.B.O.), and (*D, H*) cardiac gene expression of *Nppa, Myh7, Fibronectin, Mct2*, and *Bdh1* in mice on ND (left panel) and HFD (right panel). Data are individual mice with means ± SD for Fig *A, B, D* (*Myh7, Fibronectin, Mct2*, and *Bdh1*) or median ± IQR for Fig *D* (*Nppa*) (*n*=6-7/group). Fig *D* (*Nppa*): Statistics performed on log transformed data. * *P* < 0.001 *vs* SHAM counterpart, # *P* < 0.01 *vs* VEH counterpart by Tukey’s *post hoc* following significant interaction by two-way ANOVA. Fig *D* (*Mct2* in ND mice): no differences observed with Tukey’s *post hoc* despite significant interaction by two-way ANOVA. NS=not significant.

**Figure 3.**
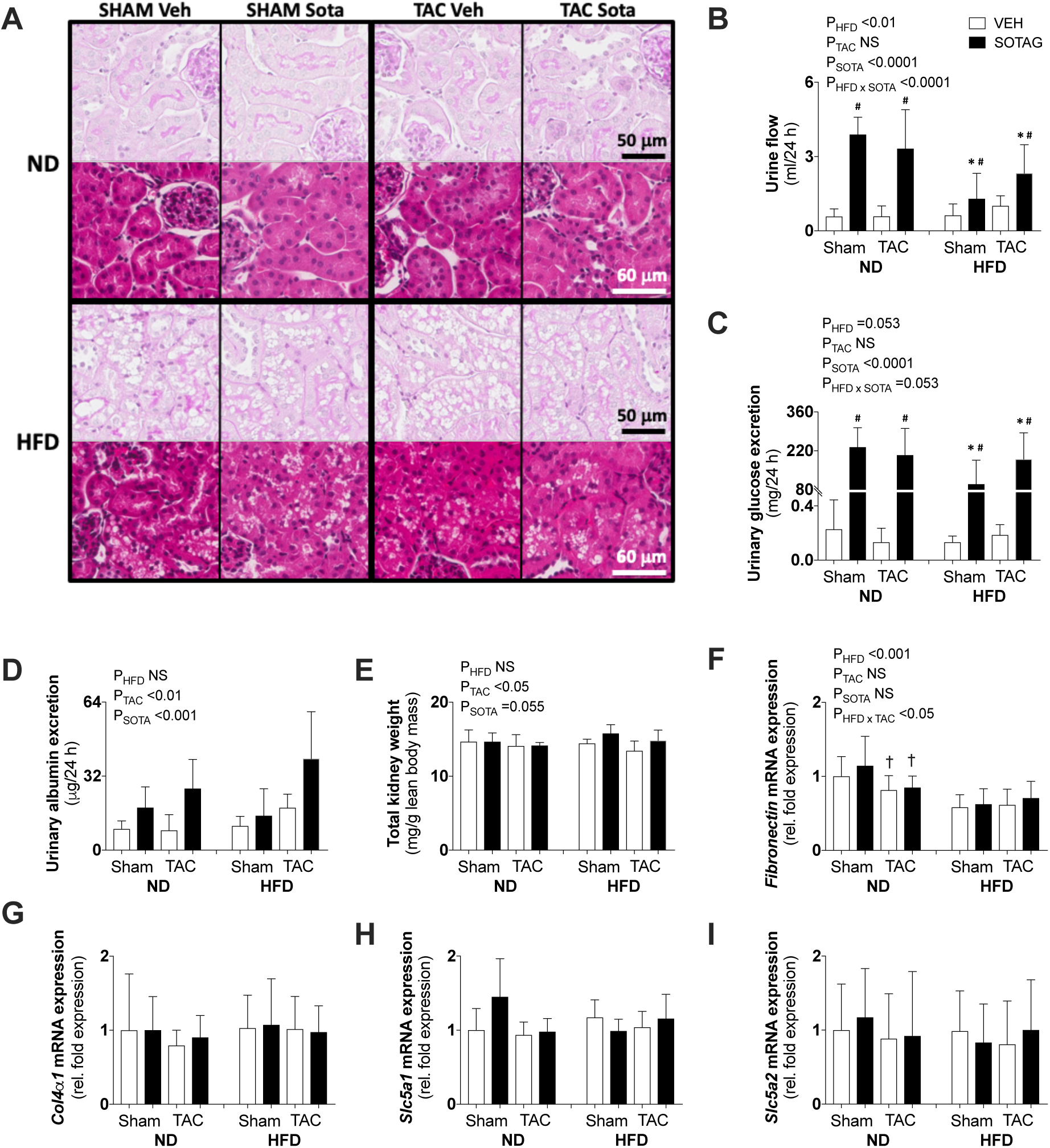
Kidney pathology in non-diabetic and diabetic mice with cardiac pressure overload treated with sotagliflozin. (*A*) Kidney histopathology representatives of periodic acid Schiff (PAS) and H&E staining used in descriptive histological assessments by expert pathologist (H.B.O.), (*B*) 24-hour urine production, (*C*) 24-hour urinary glucose excretion, (*D*) albuminuria, (*E*) kidney weight, (*F-I*) kidney gene expression of *Fibronectin, ColIVa1, Scl5a1*, and *Scl5a2*. Data are means ± SD (*n*=5-7/group). * *P* < 0.01 *vs* ND counterpart, # *P* < 0.01 *vs* VEH counterpart, † *P* < 0.05 *vs* SHAM counterpart following significant interaction by three-way ANOVA. Interaction p-values indicated only when significant (or trend). NS=not significant.

**Figure 4.**
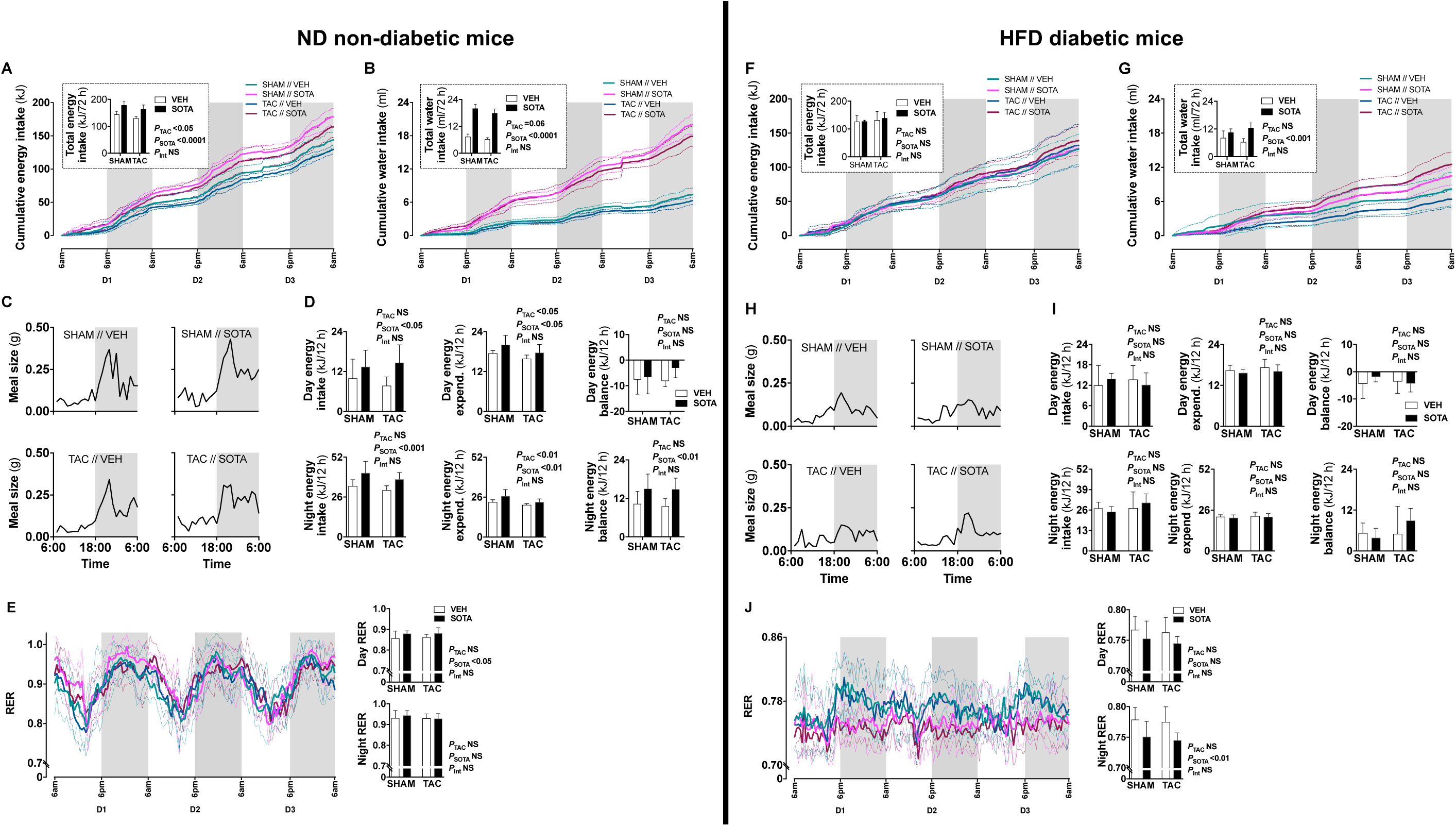
Food and water patterns, and energy balance in non-diabetic and diabetic mice with cardiac pressure overload treated with sotagliflozin. (*A, F*) Energy intake, (*B, G*) water intake, (*C, H*) eating patterns (mean meal size each hour), (*D, I*) energy balance (excluding calorie loss with urinary glucose excretion, sotagliflozin: <0.006 kJ/24 h, vehicle: <0.00001 kJ/24 h), and (*E, J*) respiratory exchange ratio (RER) in mice on ND (left panel) and HFD (right panel). Data are means ± SD denoted by dotted lines or error bars (*n*=5-7/group). NS=not significant.

### Sotagliflozin treatment lowered blood glucose and plasma insulin levels in both ND non-diabetic and HFD diabetic mice

Sotagliflozin reduced blood glucose and plasma insulin levels in both ND and HFD groups, (*Figure 1B-F*). There was no effect of sotagliflozin on body mass or body fat percentage (*Figure 1G-H*).

### Sotagliflozin treatment improved heart failure risk factors in ND non-diabetic, but not HFD diabetic, mice

In ND mice, treatment with sotagliflozin reduced TAC-induced cardiac hypertrophy, including reductions in whole heart weight, cardiomyocyte size, and left ventricular anterior and posterior wall thickness (*Figure 2A-B* and *Table 1*). Histological assessment revealed that TAC-induced pathology was mostly absent in ND mice treated with sotagliflozin. These structural benefits were not associated with a significant reduction in cardiac *Myh7* gene expression nor any alteration to other genes involved in fibrosis (*Nppa* and *Fibronectin*) or ketolysis (*Mct2* and *Bdh1, Figure 2D*). On the other hand, in HFD mice, sotagliflozin did not improve TAC-induced cardiac hypertrophy, histopathology, nor increased cardiac gene expression of *Mhy7, Nppa*, and *Fibronectin* (*Figure 2E-H* and *Table 2*). In fact, treatment with sotagliflozin in HFD TAC- and sham-operated mice increased relative gene expression of *Nppa* compared with vehicle (*Figure 2H*). In HFD mice, sotagliflozin did not attenuate the TAC-induced decrease in *Mct2* nor increased *Bdh1* (*Figure 2H*).

### Reduced efficacy of sotagliflozin in HFD diabetic mice is associated with proximal tubule injury

In all HFD mice, histopathological assessment of PAS- and Trichrome-stained slides revealed micro- and macrovacuolation of proximal tubular cells, largely restricted to segments 1 and 2 (S1/2) (*Figure 3A*). H&E staining did not reveal protein globules, implicating the vacuolation as being due to fat accumulation with mild cytoplasmic/membranous admixture in these HFD kidneys. The vast majority of glomeruli were considered normal. There was no kidney pathology induced or exacerbated by TAC or improved with sotagliflozin treatment. Associated with this kidney pathology was a lessened diuretic and glucosuric effect of sotagliflozin in HFD *vs* ND mice (*Figure 3B-C*). HFD mice did not exhibit changes in albuminuria or kidney weight but had reduced kidney gene expression of *Fibronectin* (*Figure 3D-F*).

TAC surgery and sotagliflozin treatment both increased albuminuria (*Figure 3D*). Kidney weight was reduced in all TAC mice, while sotagliflozin tended to increase in kidney weight (*Figure 3E*). In ND mice, there was a TAC-induced reduction in *Fibronectin* gene expression compared with SHAM (*Figure 3F*). Kidney mRNA expression of *Col4a1, Slc5a1*, and *Slc5a2* was not affected by HFD, TAC, or sotagliflozin treatment (*Figure 3G-I*).

### Sotagliflozin promoted increased energy intake and positive energy balance in ND non-diabetic, but not HFD diabetic mice

In ND mice, treatment with sotagliflozin increased energy and water intake, with more ‘grazing’ night-time eating patterns compared with vehicle (*Figure 4A-C*). This was associated with a smaller increase in energy expenditure (*i.e.* diet-induced thermogenesis), resulting in positive energy balance during night hours (*Figure 4D*). Adjustments for calorie loss via glucosuria (sotagliflozin mice: <0.006 kJ/24h, vehicle mice: <0.00001 kJ/24 h) did not appreciably lower this positive energy balance in ND mice (data not shown). Increased day energy intake was associated with increased respiratory exchange ratio (RER), reflecting increased carbohydrate utilization, compared with vehicle-treated mice (*Figure 4E*). In HFD mice, sotagliflozin had no effect on energy intake or eating patterns but resulted in increased total water intake (*Figure 4F-H)*. There was no effect of sotagliflozin on energy expenditure nor energy balance in HFD mice, although night-time RER was reduced, reflecting increased utilization of fat (*Figure 4I-J*). These changes to eating, drinking and/or overall metabolism were seen equally in SHAM and TAC animals.

## Discussion

This study examined the cardio-renal effects of the novel glucose lowering agent, sotagliflozin, in a previously characterised murine model of cardiac pressure overload with and without diabetes (15). In ND non-diabetic mice, treatment with sotagliflozin attenuated cardiac hypertrophy and histological markers of cardiac fibrosis that were induced by mechanical pressure overload. This is the first study to demonstrate a cardio-protective effect with a dual SGLT1/2 inhibitor and, importantly, argues against a negative effect of cardiac SGLT1 inhibition in the context of heart failure risk factors. These remarkable benefits, which were not seen in HFD diabetic animals, were associated with effective diuresis and glucosuria and not whole-body shifts to fatty acid utilisation. These findings demonstrate the utility of dual SGLT1/2 inhibition in treating heart failure risk factors in the non-diabetic state and provide some mechanistic insight.

Dual inhibitors are under investigation for the treatment of diabetes, and expected to yield greater glucose lowering and pleiotropic benefits compared with selective SGLT2 inhibitors (8, 9). While the cardio-protective effects of selective SGLT2 inhibitors are well established in patients with type 2 DM (1-3), and recently observed in both diabetic and non-diabetic patients with established heart failure in the dapagliflozin-heart failure (DAPA-HF) trial (4, 5), the efficacy of dual SGLT1/2 inhibition in this capacity is not known. In this study, TAC surgery induced a 50% increase in cardiac pressure overload in both ND and HFD mice, associated with left ventricular hypertrophic remodeling, fibrosis, leukocyte infiltration, albeit preserved ejection fraction. Sotagliflozin treatment for seven weeks overcame these mechanically induced risk factors for heart failure in ND non-diabetic mice. In a previous study using non-diabetic TAC mice with reduced ejection fraction, just two weeks’ treatment with the selective SGLT2 inhibitor, empagliflozin, attenuated a progressive decline in cardiac function (16). The authors reported no amelioration of cardiac fibrosis; however, this was not thoroughly examined, and the study lacked a sham-operated control group for reference.

The mechanisms underlying cardiac benefits with SGLT inhibition remain unclear but are likely to involve both direct and indirect effects. There is particular interest around changes to myocardial energetics, induced by direct actions to cardiac machinery as well as systemic changes to substrate availability. In humans with and without type 2 DM, empagliflozin decreased RER indicating increased fatty acid utilisation (17). Reduced insulin levels with SGLT inhibition, as seen in this study, increases lipolysis and circulating free fatty acid levels. This, in turn, stimulates hepatic ketogenesis and increases circulating ketone levels as reported in both non-diabetic and diabetic subjects treated with SGLT2 inhibitors (17, 18). With respect to the failing heart, evidence from both humans and animals show that it becomes increasingly reliant on ketone bodies as a source of fuel, due to impaired free fatty acid and glucose oxidation (19). Therefore, increased ketone availability under SGLT blockade may ‘rescue’ the failing heart, translating into cardiovascular benefits. In a non-diabetic porcine model of heart failure, treatment with empagliflozin partially restored myocardial free fatty acid uptake and increased myocardial ketone body uptake by four-fold (20). This was associated with increased myocardial activity and/or expression of ketone oxidation enzymes and ATP content as well as improved cardiac structure and contractile function. In non-diabetic rats with surgically-induced myocardial infarction, empagliflozin treatment commencing either prior to, or after, surgery, similarly increased circulating ketone levels and myocardial ketone utilisation (increased enzyme expression), associated with ameliorated cardiac hypertrophy, interstitial fibrosis, and oxidative stress and improved left ventricular function (21).

In the current study, we observed no sotagliflozin-induced changes to cardiac expression of ketolysis-associated genes but, rather, a shift toward whole-body carbohydrate utilisation and positive energy balance in non-diabetic mice on standard chow. The latter were mediated by increased energy intake with sotagliflozin and, hence, continuous carbohydrate fuel availability. Increased energy intake is seen in both humans and animals treated with SGLT inhibitors, as an adaptive response to glucosuria-induced energy loss (22). In our high fat-fed mice, however, there was no sotagliflozin-induced increase in energy intake and night-time RER decreased even further, beyond that caused by high fat feeding alone. Hence, a shift toward whole-body fatty acid utilization with SGLT inhibition appears to depend on a restriction in carbohydrate availability and may not be critical for the induction of cardio-protective effects. In diet-induced obese rats treated with dapagliflozin, pair-feeding to match food intake of vehicle-treated animals resulted in a four-fold greater weight loss (23). Therefore, additional cardiac benefits may become apparent with restricted food availability and a shift toward neutral or mild negative energy balance (irrespective of weight loss), which requires further study. It is worth noting that we did not examine fatty acid or glucose oxidation at the level of the heart *per se*; and this may have differed from whole-body RER measurements.

In the current study, cardiac benefits with sotagliflozin, seen in ND non-diabetic mice, were not evident in the HFD diabetic group, which provides additional mechanistic insight. A key difference between the two groups was the reduced efficacy with respect to sotagliflozin-induced diuresis and glucosuria in HFD animals. While treatment with sotagliflozin did lower blood glucose levels in HFD mice, these were not normalized to ND levels and the resultant diuresis and glucosuria were only ∼50-65% of that in ND mice. This was associated with compromised kidney morphology in HFD mice, largely restricted to the renal site of drug action, *i.e.*, the proximal tubule, which was not attenuated with sotagliflozin treatment. Fat deposition in proximal tubules is consistent with our previous observations in this model, where we also reported increased urinary excretion of the proximal tubule damage marker, KIM-1 (15). Others have also reported impaired sodium handling in high fat fed mice, suggestive of impaired proximal tubular function (24). Hence, the lack of cardiovascular benefit with sotagliflozin in our high fat fed TAC animals may arise from ineffective proximal tubular drug action and/or diuresis and natriuresis, otherwise thought to improve cardiovascular hemodynamics, including reduced cardiac preload (25). In the EMPA-REG trial, carried out in patients with type 2 DM and preserved renal function, 50% of the cardiovascular benefit was attributed to increased haematocrit (26). While this may reflect volume contraction, increased erythropoiesis has also been implicated; and this too can exert cardio-protective effects (27). As a dual inhibitor, sotagliflozin also mediates its blood glucose lowering effects at the level of small intestine, by delaying glucose absorption and reducing postprandial glucose (28). While it is probable that the intestinal mucosa of HFD animals in this study was compromised, sotagliflozin remained effective at slowing glucose absorption during the OGTT. Moreover, given the glucose-independent cardiovascular benefits with selective SGLT2 inhibitors (4, 5), it is unlikely that reductions in post-prandial glucose in ND mice mediated the cardiac benefits. However, we cannot rule out other cardio-protective mechanisms from inhibited intestinal glucose absorption, such as increased glucagon-like peptide 1, which may have been compromised in HFD mice.

Alternative mechanisms proposed for cardio-protection, for selective SGLT2 inhibitors at least, include other kidney effects that subsequently benefit heart function (e.g. changes in the renin angiotensin aldosterone system), inhibition of the sympathetic nervous system, and/or reduced cardiac cytosolic Na^+^ and restored Ca^2+^ handling through direct cardiac NHE-1 inhibition. A recent study in non-diabetic rodents, with either reduced or preserved ejection fraction, found that cardio-protection with empagliflozin was not associated with changes in cardiac ketone oxidation but, rather, reduced cardiac inflammation through attenuated activation of the nucleotide-binding domain-like receptor protein 3 (NLRP3) inflammasome, which was dependent on restoration of cardiac cytoplasmic Ca^2+^ levels (29). Any additional actions of dual SGLT1/2 inhibition, due to the cardiac presence of SGLT1, remain to be elucidated.

Heart failure is one the most common cardiovascular manifestations in patients with type 2 DM (30). While our findings suggest that sotagliflozin was not effective for the treatment of heart failure risk factors in the obesogenic-diabetic state, it is important to note that in the DAPA-HF trial, both obese and type 2 DM patients equally benefitted with respect to cardiovascular outcomes (4, 5). It is possible that our high fat fed mice, with severely compromised renal morphology, do not reflect the kidney status of the trial patients. Nevertheless, our results indicate that the efficacy of this agent in treating heart failure risk factors appears to be limited in settings of proximal tubular compromise.

This study demonstrated cardio-protective effects of dual SGLT1/2 inhibition in non-diabetic, but not high fat-induced diabetic, mice with cardiac pressure overload. Our findings suggest that cardiac benefits occur in the absence of whole-body or cardiac-specific shifts toward fatty acid and/or ketone body utilisation but, rather, may be associated with profound diuresis and/or glucosuria, which was not seen in our high fat-fed diabetic mice.

## Acknowledgements

The authors wish to thank the School of Biomedical Sciences, The University of Queensland, Mater Research Institute-University of Queensland, and the Mater Foundation for institutional support.

## References

1. Kosiborod AM, Cavender ZM, Fu PA, Wilding WJ, Khunti IK, Holl ER, Norhammar EA, Birkeland EK, Jørgensen EM, Thuresson EM, Arya EN, Bodegård EJ, Hammar EN, Fenici EP. Lower Risk of Heart Failure and Death in Patients Initiated on Sodium-Glucose Cotransporter-2 Inhibitors Versus Other Glucose-Lowering Drugs: The CVD-REAL Study (Comparative Effectiveness of Cardiovascular Outcomes in New Users of Sodium-Glucose Cotransporter-2 Inhibitors). Circulation 2017; 136:249–59.

2. Fitchett D, Inzucchi SE, Cannon CP, McGuire DK, Scirica BM, Johansen OE, Sambevski S, Kaspers S, Pfarr E, George JT, Zinman B. Empagliflozin Reduced Mortality and Hospitalization for Heart Failure Across the Spectrum of Cardiovascular Risk in the EMPA-REG OUTCOME Trial. Circulation 2019; 139:1384–95.

3. Wiviott SD, Raz I, Bonaca MP, Mosenzon O, Kato ET, Cahn A, Silverman MG, Zelniker TA, Kuder JF, Murphy SA, Bhatt DL, Leiter LA, McGuire DK, Wilding JPH, Ruff CT, Gause-Nilsson IAM, Fredriksson M, Johansson PA, Langkilde AM, Sabatine MS, Investigators D-T. Dapagliflozin and Cardiovascular Outcomes in Type 2 Diabetes. N Engl J Med 2019; 380:347–57.

4. McMurray JJV, Solomon SD, Inzucchi SE, Kober L, Kosiborod MN, Martinez FA, Ponikowski P, Sabatine MS, Anand IS, Belohlavek J, Böhm M, Chiang C-E, Chopra VK, de Boer RA, Desai AS, Diez M, Drozdz J, Dukát A, Ge J, Howlett JG, Katova T, Kitakaze M, Ljungman C, Merkely B, Nicolau JC, O’Meara E, Petrie MC, Vinh PN, Schou M, Tereshchenko S, Verma S, Held C, DeMets DL, Docherty KF, Jhund PS, Bengtsson O, Sjöstrand M, Langkilde AM. Dapagliflozin in Patients with Heart Failure and Reduced Ejection Fraction. New England Journal Of Medicine 2019.

5. Petrie MC, Verma S, Docherty KF, Inzucchi SE, Anand I, Belohlavek J, Bohm M, Chiang CE, Chopra VK, de Boer RA, Desai AS, Diez M, Drozdz J, Dukat A, Ge J, Howlett J, Katova T, Kitakaze M, Ljungman CEA, Merkely B, Nicolau JC, O’Meara E, Vinh PN, Schou M, Tereshchenko S, Kober L, Kosiborod MN, Langkilde AM, Martinez FA, Ponikowski P, Sabatine MS, Sjostrand M, Solomon SD, Johanson P, Greasley PJ, Boulton D, Bengtsson O, Jhund PS, McMurray JJV. Effect of Dapagliflozin on Worsening Heart Failure and Cardiovascular Death in Patients With Heart Failure With and Without Diabetes. JAMA 2020.

6. Gallo LA, Wright EM, Vallon V. Probing SGLT2 as a therapeutic target for diabetes: Basic physiology and consequences. Diabetes & Vascular Disease Research 2015; 12:78–89.

7. Rieg T, Masuda T, Gerasimova M, Mayoux E, Platt K, Powell DR, Thomson SC, Koepsell H, Vallon V. Increase in SGLT1-mediated transport explains renal glucose reabsorption during genetic and pharmacological SGLT2 inhibition in euglycemia. American journal of physiology Renal physiology 2014; 306:F188–F93.

8. Rosenstock J, Cefalu WT, Lapuerta P, Zambrowicz B, Ogbaa I, Banks P, Sands A. Greater dose-ranging effects on A1C levels than on glucosuria with LX4211, a dual inhibitor of SGLT1 and SGLT2, in patients with type 2 diabetes on metformin monotherapy. Diabetes care 2015; 38:431.

9. Zambrowicz B, Freiman J, Brown P, Frazier KS, Turnage A, Bronner J, Ruff D, Shadoan M, Banks P, Mseeh F, Rawlins D, Goodwin NC, Mabon R, Harrison BA, Wilson A, Sands A, Powell D. LX4211, a Dual SGLT1/SGLT2 Inhibitor, Improved Glycemic Control in Patients With Type 2 Diabetes in a Randomized, Placebo-Controlled Trial. Clin Pharmacol Ther 2012; 92:158–69.

10. Kashiwagi Y, Nagoshi T, Yoshino T, Tanaka TD, Ito K, Harada T, Takahashi H, Ikegami M, Anzawa R, Yoshimura M. Expression of SGLT1 in Human Hearts and Impairment of Cardiac Glucose Uptake by Phlorizin during Ischemia-Reperfusion Injury in Mice. PLoS ONE 2015; 10:e0130605.

11. Banerjee S, McGaffin K, Pastor-Soler N, Ahmad F. SGLT1 is a novel cardiac glucose transporter that is perturbed in disease states. Cardiovascular Research 2009; 84:111–8.

12. Vrhovac I, Balen Eror D, Klessen D, Burger C, Breljak D, Kraus O, Radovic N, Jadrijevic S, Aleksic I, Walles T, Sauvant C, Sabolic I, Koepsell H. Localizations of Na(+)-D-glucose cotransporters SGLT1 and SGLT2 in human kidney and of SGLT1 in human small intestine, liver, lung, and heart. Pflugers Arch 2015; 467:1881–98.

13. Banerjee SK, Wang DW, Alzamora R, Huang XN, Pastor-Soler NM, Hallows KR, McGaffin KR, Ahmad F. SGLT1, a novel cardiac glucose transporter, mediates increased glucose uptake in PRKAG2 cardiomyopathy. J Mol Cell Cardiol 2010; 49:683–92.

14. Ramratnam M, Sharma RK, D’ Auria S, Lee SJ, Wang D, Huang XYN, Ahmad F. Transgenic Knockdown of Cardiac Sodium/Glucose Cotransporter 1 (SGLT1) Attenuates PRKAG2 Cardiomyopathy, Whereas Transgenic Overexpression of Cardiac SGLT1 Causes Pathologic Hypertrophy and Dysfunction in Mice. Journal of the American Heart Association 2014; 3:n/a-n/a.

15. Tan WS, Mullins TP, Flint M, Walton SL, Bielefeldt-Ohmann H, Carter DA, Gandhi MR, McDonald HR, Li J, Moritz KM, Reichelt ME, Gallo LA. Modeling heart failure risk in diabetes and kidney disease: limitations and potential applications of transverse aortic constriction in high-fat-fed mice. American journal of physiology Regulatory, integrative and comparative physiology 2018; 314:R858–R69.

16. Byrne NJ, Parajuli N, Levasseur JL, Boisvenue J, Beker DL, Masson G, Fedak PWM, Verma S, Dyck JRB. Empagliflozin Prevents Worsening of Cardiac Function in an Experimental Model of Pressure Overload-Induced Heart Failure. JACC Basic Transl Sci 2017; 2:347–54.

17. Ferrannini E, Baldi S, Frascerra S, Astiarraga B, Heise T, Bizzotto R, Mari A, Pieber TR, Muscelli E. Shift to Fatty Substrate Utilization in Response to Sodium-Glucose Cotransporter 2 Inhibition in Subjects Without Diabetes and Patients With Type 2 Diabetes. Diabetes 2016; 65:1190–5.

18. Polidori D, Iijima H, Goda M, Maruyama N, Inagaki N, Crawford PA. Intra- and inter-subject variability for increases in serum ketone bodies in patients with type 2 diabetes treated with the sodium glucose co-transporter 2 inhibitor canagliflozin. Diabetes Obes Metab 2018; 20:1321–6.

19. Aubert G, Martin OJ, Horton JL, Lai L, Vega RB, Leone TC, Koves T, Gardell SJ, Kruger M, Hoppel CL, Lewandowski ED, Crawford PA, Muoio DM, Kelly DP. The Failing Heart Relies on Ketone Bodies as a Fuel. Circulation 2016; 133:698–705.

20. Santos-Gallego CG, Requena-Ibanez JA, San Antonio R, Ishikawa K, Watanabe S, Picatoste B, Flores E, Garcia-Ropero A, Sanz J, Hajjar RJ, Fuster V, Badimon JJ. Empagliflozin Ameliorates Adverse Left Ventricular Remodeling in Nondiabetic Heart Failure by Enhancing Myocardial Energetics. J Am Coll Cardiol 2019; 73:1931–44.

21. Yurista SR, Silljé HHW, Oberdorf-Maass SU, Schouten E-M, Pavez Giani MG, Hillebrands J-L, van Goor H, van Veldhuisen DJ, de Boer RA, Westenbrink BD. Sodium– glucose co-transporter 2 inhibition with empagliflozin improves cardiac function in non-diabetic rats with left ventricular dysfunction after myocardial infarction. 2019; 21:862–73.

22. Ferrannini G, Hach T, Crowe S, Sanghvi A, Hall KD, Ferrannini E. Energy Balance After Sodium-Glucose Cotransporter 2 Inhibition. Diabetes Care 2015; 38:1730–5.

23. Devenny JJ, Godonis HE, Harvey SJ, Rooney S, Cullen MJ, Pelleymounter MA. Weight loss induced by chronic dapagliflozin treatment is attenuated by compensatory hyperphagia in diet-induced obese (DIO) rats. Obesity (Silver Spring) 2012; 20:1645–52.

24. Deji N, Kume S, Araki S, Soumura M, Sugimoto T, Isshiki K, Chin-Kanasaki M, Sakaguchi M, Koya D, Haneda M, Kashiwagi A, Uzu T. Structural and functional changes in the kidneys of high-fat diet-induced obese mice. Am J Physiol Renal Physiol 2009; 296:F118–26.

25. Verma S, McMurray JJV. SGLT2 inhibitors and mechanisms of cardiovascular benefit: a state-of-the-art review. Diabetologia 2018; 61:2108–17.

26. Inzucchi SE, Zinman B, Fitchett D, Wanner C, Ferrannini E, Schumacher M, Schmoor C, Ohneberg K, Johansen OE, George JT, Hantel S, Bluhmki E, Lachin JM. How Does Empagliflozin Reduce Cardiovascular Mortality? Insights From a Mediation Analysis of the EMPA-REG OUTCOME Trial. Diabetes Care 2018; 41:356–63.

27. Mazer CD, Hare GMT, Connelly PW, Gilbert RE, Shehata N, Quan A, Teoh H, Leiter LA, Zinman B, Juni P, Zuo F, Mistry N, Thorpe KE, Goldenberg RM, Yan AT, Connelly KA, Verma S. Effect of Empagliflozin on Erythropoietin Levels, Iron Stores, and Red Blood Cell Morphology in Patients With Type 2 Diabetes Mellitus and Coronary Artery Disease. Circulation 2020; 141:704–7.

28. Zambrowicz B, Lapuerta P, Strumph P, Banks P, Wilson A, Ogbaa I, Sands A, Powell D. LX4211 therapy reduces postprandial glucose levels in patients with type 2 diabetes mellitus and renal impairment despite low urinary glucose excretion. Clin Ther 2015; 37:71–82 e12.

29. Byrne NJ, Matsumura N, Maayah ZH, Ferdaoussi M, Takahara S, Darwesh AM, Levasseur JL, Jahng JWS, Vos D, Parajuli N, El-Kadi AOS, Braam B, Young ME, Verma S, Light PE, Sweeney G, Seubert JM, Dyck JRB. Empagliflozin Blunts Worsening Cardiac Dysfunction Associated With Reduced NLRP3 (Nucleotide-Binding Domain-Like Receptor Protein 3) Inflammasome Activation in Heart Failure. Circ Heart Fail 2020; 13:e006277.

30. Shah AD, Langenberg C, Rapsomaniki E, Denaxas S, Pujades-Rodriguez M, Gale CP, Deanfield J, Smeeth L, Timmis A, Hemingway H. Type 2 diabetes and incidence of cardiovascular diseases: a cohort study in 1.9 million people. Lancet Diabetes Endocrinol 2015; 3:105–13.

